# AI-Based Detection of Coliform Colonies Using CNN Transfer Learning for Application to Cultured Plate Analysis in Water Quality Research

**DOI:** 10.1101/2025.06.29.662231

**Authors:** Shivaji Mallela, Abria Gates, Sandeep Medepalli, Benedict Okeke, Olcay Kursun

## Abstract

Pathogenic bacterial contamination of water poses a severe public health risk, particularly in settings with limited laboratory resources. We propose a two-stage artificial intelligence (AI) pipeline for automated detection and classification of coliform colonies on agar plates. In the first stage, a YOLOv8-based detector localizes colonies on full-plate images, eliminating the need for manual annotation. In the second stage, detected colony patches are classified using a convolutional neural network (CNN) trained via transfer learning, where models are first pretrained on a diverse public bacterial colony dataset and subsequently fine-tuned on coliform-specific classification tasks. Across both in-house and public datasets, transfer learning consistently improves classification performance relative to training from scratch. The complete pipeline processes each plate in under five seconds and outperforms classical feature-based baselines, including Histogram of Oriented Gradients, Local Binary Patterns, and Haralick descriptors with conventional classifiers. These results demonstrate the potential of a modular, low-cost AI framework for scalable and accessible microbiological analysis, with future work targeting color-aware models and on-device inference for field deployment.

## 1. Introduction

Waterborne diseases remain a major public health challenge worldwide, with an estimated 1.6 million deaths annually linked to unsafe water and sanitation [1]. Fecal contamination, often stemming from agricultural runoff, sewage leaks, or inadequate sanitation, introduces pathogenic microorganisms into drinking and recreational water supplies. Among these, coliform bacteria such as *Escherichia coli* serve as reliable sentinel organisms: their presence correlates strongly with fecal pollution and the potential co-occurrence of more dangerous pathogens like *Salmonella* species or *Vibrio* cholerae [2,3].

Conventional microbiological assays for coliform colony localization and classification involve sample filtration or plating on selective agar media, followed by 24-48 hours of incubation and manual colony enumeration. Though well established, these workflows are time-consuming, labor-intensive, and require specialized laboratory facilities and trained personnel [4–6]. In remote or resource-limited settings, such constraints often delay critical interventions, sometimes by days, undermining efforts to prevent outbreaks and protect vulnerable communities [6]. Recent efforts to automate bacterial colony counting have explored deep learning approaches [7–9],some recent studies have applied deep learning to bacterial colony detection and analysis on agar plates, often using two-stage detection-classification pipelines or CNN-based counting approaches, including TFT sensor-based colony analysis [23], CNN-based microbial colony detection [24], deep learning for vaccine production monitoring [25], and digital microbiology imaging methods [26]. Our pipeline is conceptually aligned with these approaches but differs in its explicit decoupling of colony localization and classification using a modern YOLO-based detector followed by CNN classifiers with transfer learning. These subtle differences allow a scalable approach that does not require large training datasets, does not rely solely on color-specific cues, or lack adaptability across diverse field conditions. Commercial automated colony counters (e.g., ProtoCOL [27], ScanStation [28]) are reliable in controlled settings but are typically proprietary, cost-intensive, and limited in flexibility. In contrast, our software-based pipeline is modular and adaptable, enabling retraining across species, imaging conditions, and devices, while also serving as a platform for training students to develop and extend new analysis tools as needed. To address these issues, our interdisciplinary team supported by the NSF ExpandAI program has developed an end-to-end, AI-powered framework for automated colony detection and classification on agar plates. Leveraging advances in computer vision and deep learning, our system replaces manual preprocessing (previously performed in ImageJ [4] or EBImage [10]) with a YOLOv8 object detector [11] that rapidly localizes colonies. A second stage employs transfer-learned convolutional neural networks (CNNs) trained on a diverse 10-class dataset [12] to differentiate coliform species with high precision. By combining these two components into a cohesive pipeline, we achieve accurate and scalable analysis in seconds per plate for low-cost, field-deployable water quality monitoring and opening new avenues for interdisciplinary research in environmental health science.

From a deployment perspective, imaging-based colony analysis is sensitive to acquisition variability arising from differences in cameras, lighting, and laboratory setups. In this work, potential bias from device-specific effects is partially mitigated by pretraining base models on a public dataset collected using diverse imaging conditions prior to fine-tuning on in-house data. We emphasize that the proposed system is intended as a decision-support tool rather than a standalone diagnostic, and that any public health deployment would require appropriate validation, calibration, and human oversight.

## 2. Materials and Methods

### 2.1 Dataset Collection and Description

In this study, we leverage two complementary image collections to develop and evaluate our two-stage AI pipeline. The Source (Pretraining) Dataset comprises a publicly available repository of high-resolution agar-plate photographs spanning 24 bacterial species [12], from which we drew ten classes (≈5,000 patches) to pretrain both our YOLOv8 detector and the initial convolutional backbone. In contrast, the Target (Fine-Tuning) Dataset consists of images we collected in-house at Auburn University at Montgomery (AUM) of four coliform species [13], which are *Citrobacter freundii*, *Enterobacter aerogenes*, *Escherichia coli*, and *Klebsiella pneumoniae*. Detection of these species is the target task that we picked for downstream fine-tuning, validation, and benchmarking of classification performance.

#### 2.1.1 In-house Dataset (Target Dataset)

Our local dataset [13] was generated by cultivating four coliform species on tryptic soy agar (TSA) plates in the laboratory. Cell suspension of C*itrobacter freundii* (OD_600_ = 0.062), *Enterobacter aerogenes* (OD_600_ = 0.034), *Escherichia coli* (OD_600_ = 0.064) and *Klebsiella pneumoniae* (OD_600_ = 0.058) were serially diluted (10^-1^ to 10^-9^) and plated on TSA plates. TSA was selected as a non-differential medium to provide a controlled morphological baseline, focusing learning on colony shape and texture rather than medium-specific coloration (extension to selective or chromogenic media will be explored as future work). Cultures were incubated at 37°C for 48 hours. Once incubation was complete, photographs of discrete colonies on each agar plate were acquired using a modern smartphone camera. The use of a cellphone streamlined image acquisition and demonstrated the feasibility of low-cost, field-friendly data collection as an important consideration for potential deployment in resource-limited settings.

For our initial classification experiments, colony detection was performed manually in ImageJ. We annotated each of the four plate images (Figure 1) to draw bounding boxes around individual colonies, then exported these regions as 185 discrete patches (approximately 40-50 patches per species). These manually cropped patches became the foundation of our early deep-learning trials, allowing us to benchmark simple CNN architectures against hand-crafted feature methods. By starting with a small but carefully labeled dataset, we could rapidly iterate on model design and preprocessing techniques without the overhead of large-scale annotation. The initial annotation process is later replaced by the proposed YOLO pipeline that is followed by CVAT (https://www.cvat.ai/) for manual corrections and published in [13]. To further evaluate the robustness of our pipeline, in a separate laboratory facility at AUM, we independently cultured and imaged an additional set of samples corresponding to the same four species; and, we also prepared mixed-coliform culture plates containing all four species on a single agar dish (Figure 2). This mixed plate served as a realistic test case: after applying our trained YOLO-based detector, the system successfully localized and then classified each colony patch via the transfer-learned CNN. The mixed-culture demonstration highlighted the end-to-end capability of our approach to potentially handle heterogeneous samples, exactly the scenario water-quality monitors might face in the field.

**Figure 1.**
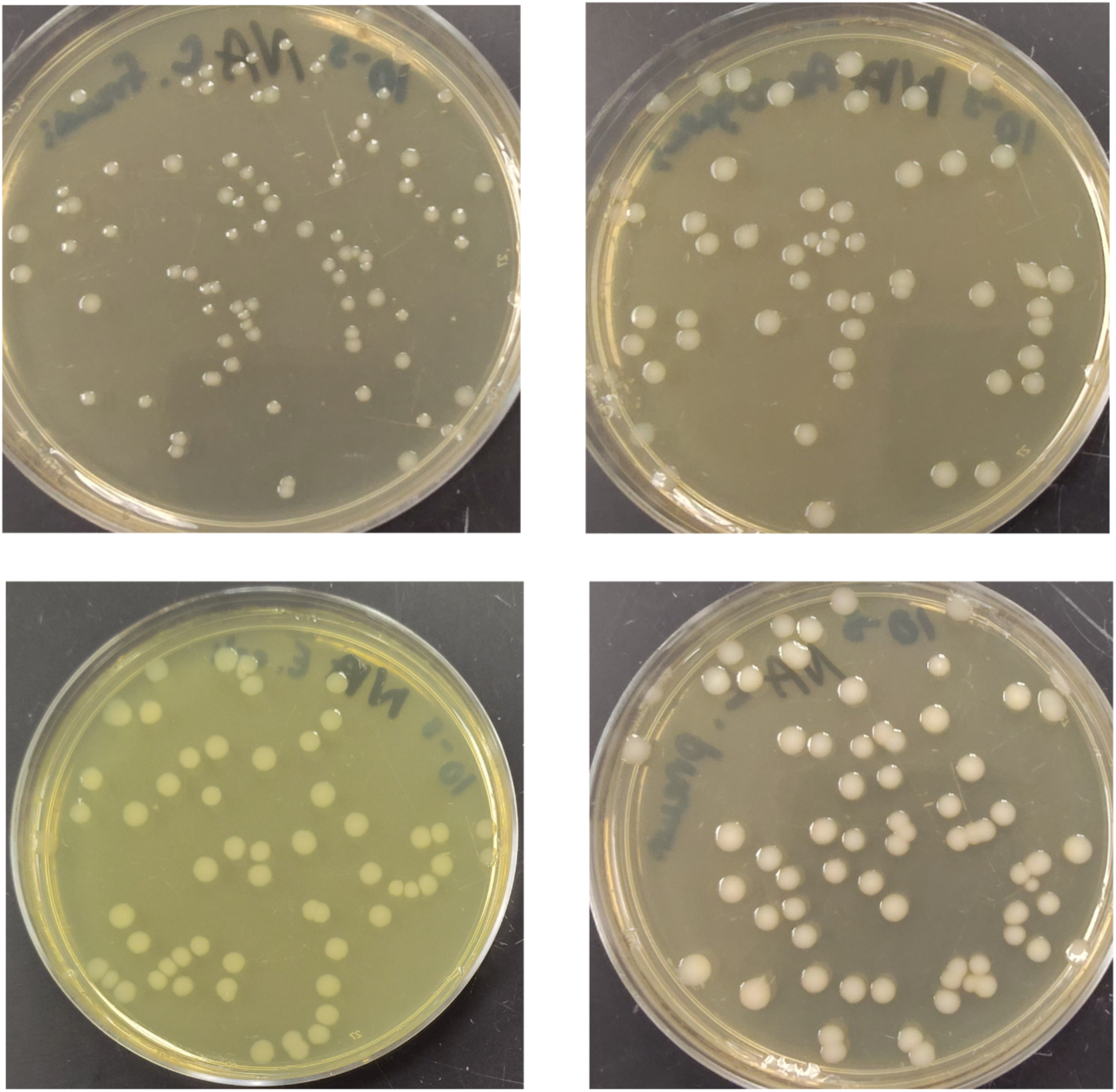
Representative images from the target (fine-tuning) dataset, showing the four coliform species in panel order from upper left to lower right: *Citrobacter freundii* (upper left), *Enterobacter aerogenes* (upper right), *Escherichia coli* (lower left), and *Klebsiella pneumoniae* (lower right).

**Figure 2.**
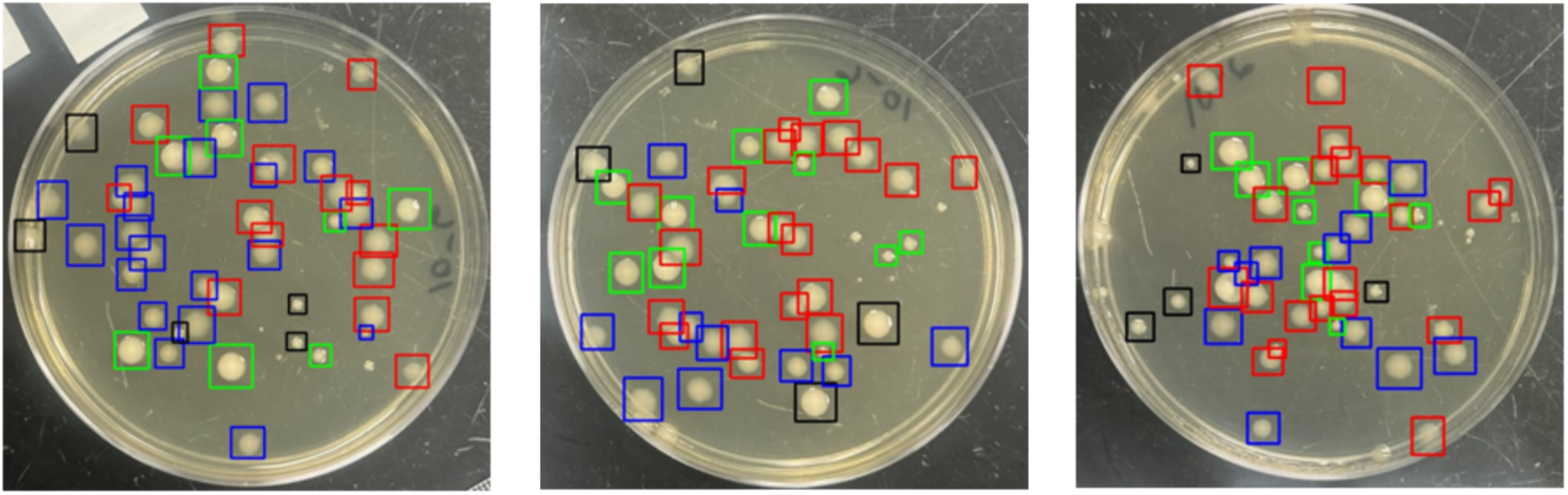
Three mixed-coliform agar plate samples from our in-house dataset, each containing colonies of *Citrobacter freundii* (Red), *Enterobacter aerogenes* (Green), *Escherichia coli* (Blue), and *Klebsiella pneumoniae* (Black). Bounding boxes overlaid on each plate indicate colony detections generated by the proposed YOLOv8 + CNN pipeline.

#### 2.1.2 Public Dataset for Transfer Learning (Pretraining dataset)

The public dataset [12] (Figure 3) comprises 369 high-resolution images of bacterial cultures on solid media, with 56,865 colonies manually annotated via bounding boxes. Images were acquired under realistic lab conditions using three different smartphone models (LG Nexus 5X, iPhone 6, Huawei P30 Lite) and both black and white backgrounds to introduce variability in lighting and device optics. Expert bacteriologists curated and validated all annotations using COCO Annotator v0.11.1, and the dataset is distributed in multiple formats (COCO JSON, Pascal VOC XML, YOLO, CSV/TSV) to facilitate diverse training workflows [12].

**Figure 3.**
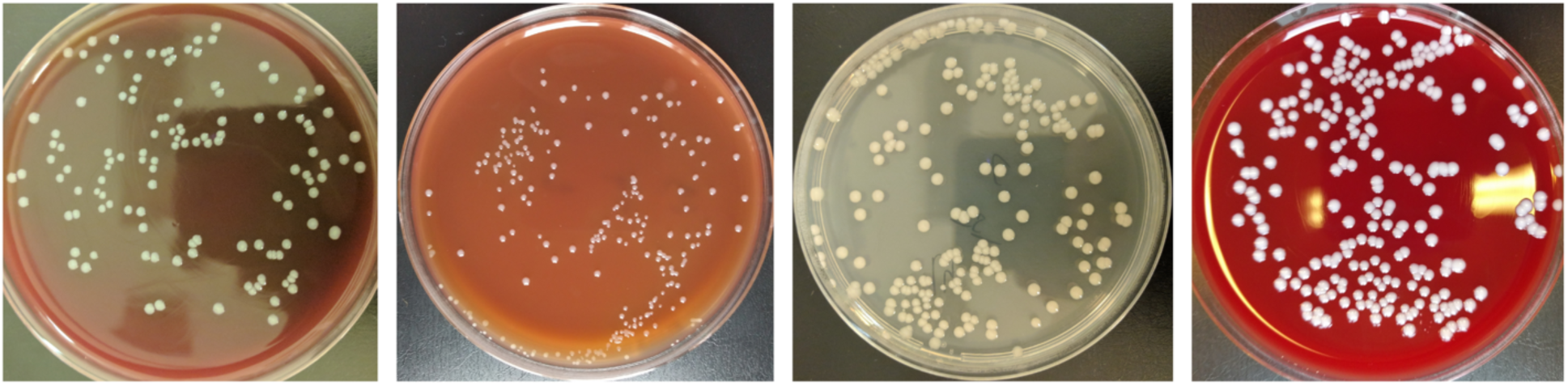
Sample images from the **Source (Pretraining) Dataset**, illustrating four of the pretraining classes: *Listeria monocytogenes*, *Pasteurella multocida*, *Salmonella enterica*, and *Staphylococcus hyicus*.

From this broad collection of 24 species, we selected 10 taxonomically distinct classes: *Actinobacillus pleuropneumoniae, Bibersteinia trehalosi, Bordetella bronchiseptica, Brucella ovis, Erysipelothrix rhusiopathiae, Glaesserella parasuis, Listeria monocytogenes, Pasteurella multocida, Rhodococcus equi, and Staphylococcus aureus*. These classes used as the base task encourages our CNN’s early layers to learn generalizable colony features rather than overfit to our four-species target set. Approximately 450 patches per class were extracted from the annotations, yielding ∼4,500 training samples for base-model pretraining.

In our pipeline, we leveraged the public dataset in three complementary ways to maximize its impact on both detection and classification performance. First, we trained the YOLOv8 detector on 119 full-plate images drawn from the public collection (YOLO performance was consistent across modest variations in class composition and image counts), collapsing all annotated colonies into a single “colony” class. Regardless of species, by targeting any colony identically during this stage, the detector learned to generalize across a wide spectrum of colony shapes, sizes, and textures, and to produce tight bounding boxes under varying lighting and background conditions. This automated annotation step replaced the manual ImageJ workflow and formed the basis for patch extraction in downstream stages.

Second, we repurposed the detector’s outputs to assemble a large, diverse training corpus for our base CNN. Every colony that YOLO localized was cropped, zero-padded, and resized to 100×100 px, yielding approximately 4,500 image patches spanning ten taxonomically distinct bacterial species. These patches served as the pretraining set for a CNN classifier, which was fine-tuned from ImageNet initialization. By learning generic colony features, rather than memorizing patterns specific to our four laboratory-cultured species, the CNN developed robust and transferrable representations that accelerated convergence. These robust features also raised validation accuracy during subsequent fine-tuning on our smaller, laboratory-collected dataset.

Finally, to demonstrate the broader applicability of our transfer-learning strategy, we conducted an additional validation using four further public classes (*Listeria monocytogenes*, *Pasteurella multocida*, *Salmonella enterica*, and *Staphylococcus hyicus*). Following the same pretrain-and- fine-tune workflow, the CNN pretrained on the original ten classes was readily adapted to these novel species, achieving comparable classification performance without architectural changes. This result underscores the versatility of our approach and its potential for rapid extension to new bacterial targets as more annotated data becomes available.

### 2.2 Proposed Method: Two-Stage Deep-Learning Pipeline

To automate and scale bacterial-colony analysis on agar plates, we designed a two-stage framework that first localizes colonies via object detection and then classifies each instance with a deep convolutional network. By chaining a YOLOv8 detector with a transfer-learned CNN, our pipeline replaces manual ImageJ annotation and handcrafted feature engineering with an end-to-end learning approach optimized for both speed and accuracy. Training was conducted on an AWS EC2 g5.xlarge instance equipped with an NVIDIA GPU, with Google Colab used as an alternative execution environment. Runtime measurements on the EC2 system showed end-to-end inference times of under five seconds per plate (with a batch size of one - without batching or throughput amortization across multiple plates). Comparable performance was observed on Google Colab using an NVIDIA T4 GPU. End-to-end runtime includes image loading, preprocessing, tiling, colony detection, patch extraction, and CNN-based classification.

#### 2.2.1 Image Preparation and Preprocessing

In this initial phase of the study, every agar-plate photograph, whether acquired in our in-house lab or drawn from public repositories, was converted to 8-bit grayscale [14]. This normalization step strips away variations in media coloration and lighting, channeling the detector’s attention toward colony morphology, texture, and shape. Although this first implementation leverages grayscale imagery to establish baseline performance, the following section extends the analysis to full-color images. In future work, the framework can be further expanded to incorporate chromogenic media and auxiliary sample metadata (e.g., source, incubation temperature, pH) through parallel network branches, providing additional contextual information to support downstream classification [15].

#### 2.2.2 Stage 1 - Colony Detection with YOLO

In the first stage of our pipeline, we leverage YOLO-Nano [11], a lightweight variant of the You Only Look Once (YOLO) architecture optimized for resource-constrained environments, to automate colony localization. YOLO is a state-of-the-art, real-time object detection algorithm that simultaneously identifies and localizes objects within an image using a single convolutional neural network. It divides the input image into a grid and, for each grid cell, predicts bounding boxes, confidence scores, and class probabilities. This architecture achieves high inference speed with competitive accuracy, making YOLO well suited for applications requiring real-time processing or limited computing resources. While YOLOv8-Nano prioritizes computational efficiency, deeper YOLO variants may provide improved localization accuracy and higher IoU at the cost of increased computational demand.

In our study, we employed YOLOv8 [11] as the colony detector. It was trained on 119 plate images from the aforementioned pretraining dataset (see Figure 4) and evaluated on a held-out test set of 105 images. All annotated colonies regardless of species were collapsed into a single “colony” label, directing YOLO to focus exclusively on precise bounding-box placement rather than species discrimination. We opted not to retrain YOLO for colony classification, as doing so would require a large number of annotated examples for each species. Thus, we used YOLO strictly as a colony detector. YOLOv8’s ability to accurately detect colonies allowed us to replace manual annotation tools such as ImageJ, significantly accelerating the preprocessing workflow for a scalable downstream classification.

**Figure 4.**
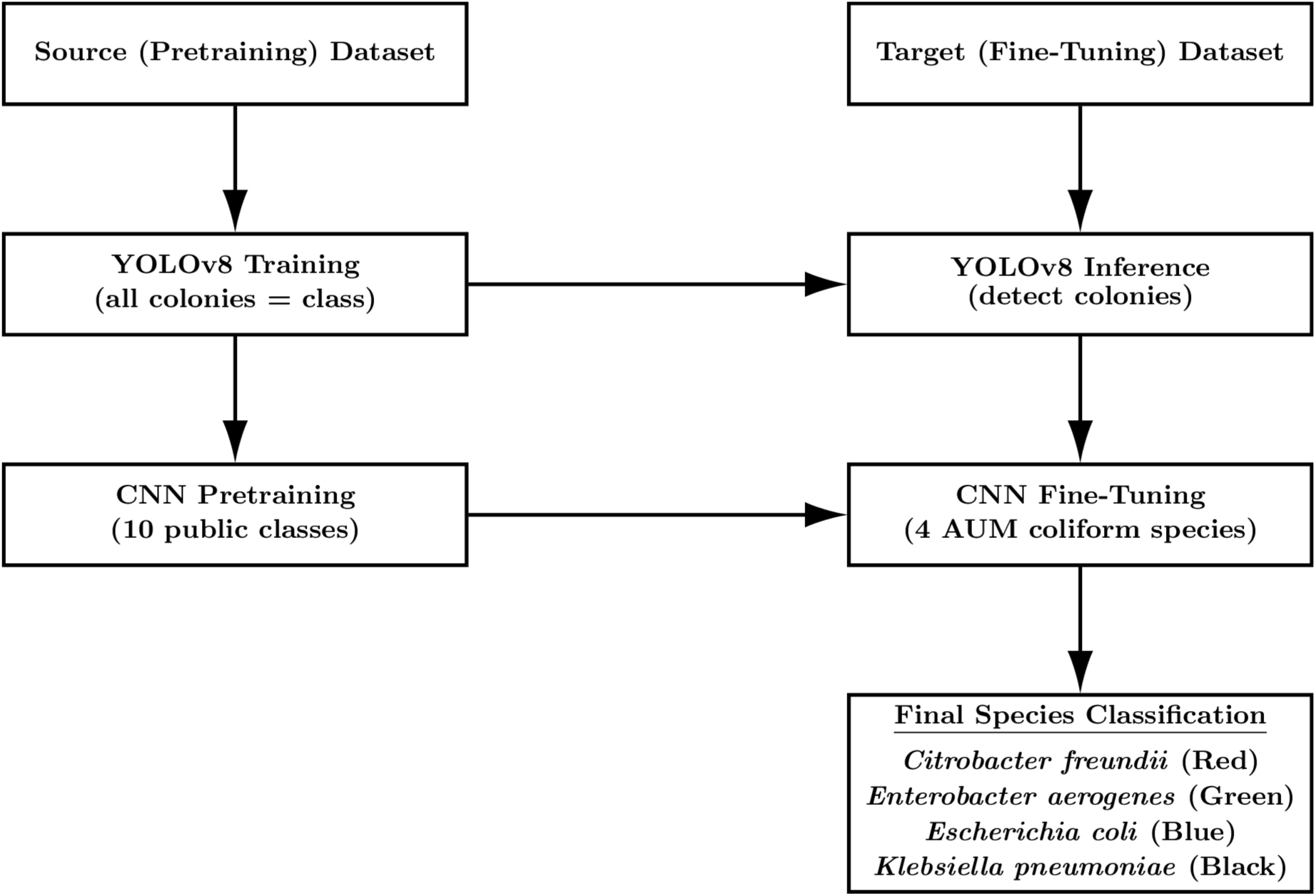
Two-column workflow illustrating the use of the Source (Pretraining) Dataset (left) and the Target (Fine-Tuning) Dataset (right). In the source domain, YOLOv8 is trained to detect colonies and a CNN is pretrained on ten public bacterial classes. These pretrained components are then transferred to the target domain, where YOLOv8 performs colony detection on in-house laboratory images and the CNN is fine-tuned on four local coliform species. The final model performs colony classification for downstream analysis.

**Figure 5.**
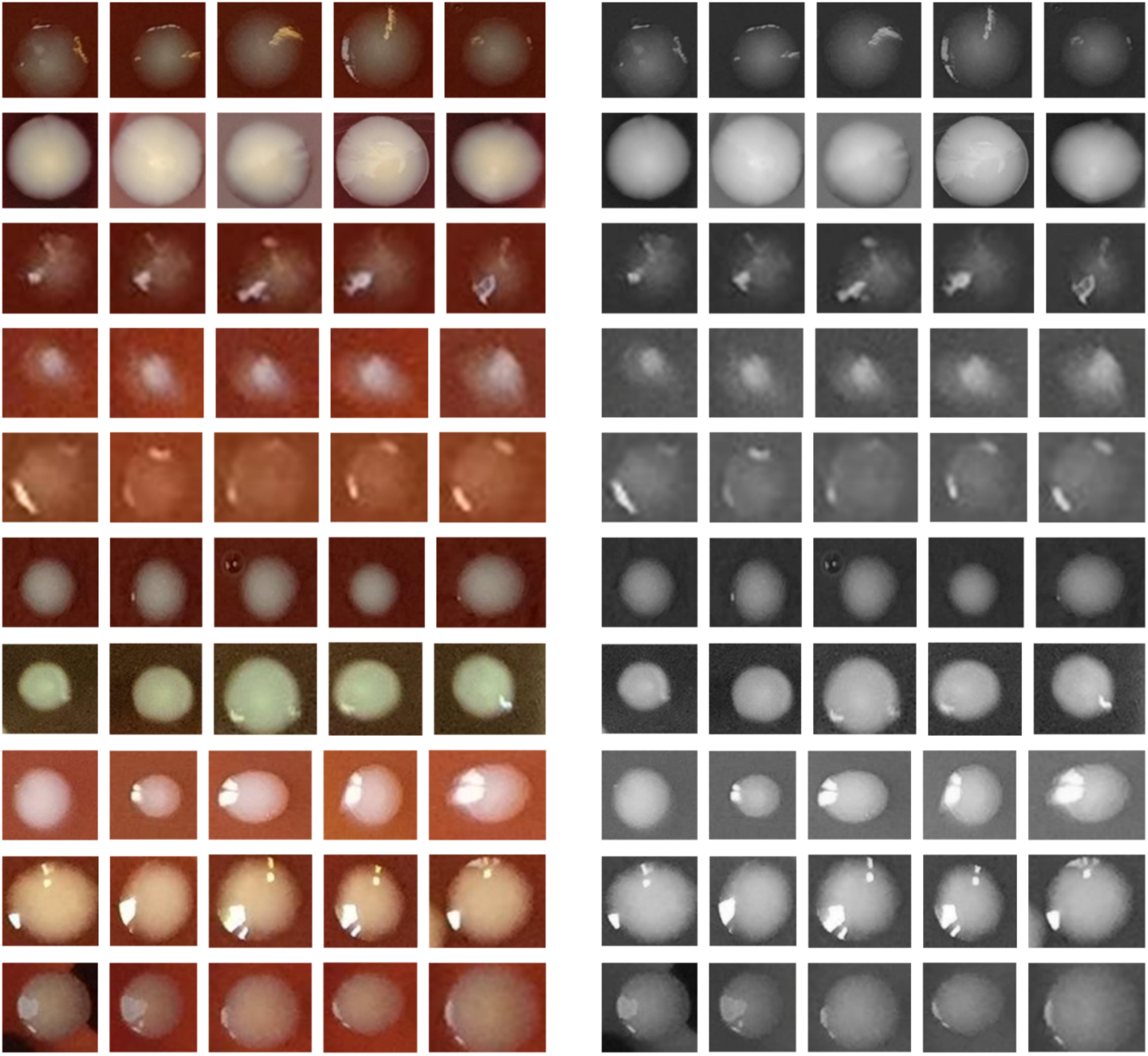
Representative samples from the ten base classes used for pretraining, arranged in a 10 × 5 grid (five patches per class). Each row corresponds to one species: sp02 *Actinobacillus pleuropneumoniae*, sp05 *Bibersteinia trehalosi*, sp06 *Bordetella bronchiseptica*, sp07 *Brucella ovis*, sp10 *Erysipelothrix rhusiopathiae*, sp12 *Glaesserella parasuis*, sp14 *Listeria monocytogenes*, sp16 *Pasteurella multocida*, sp19 *Rhodococcus equi*, and sp21 *Staphylococcus aureus*. The left panel shows original color crops and the right panel shows the corresponding 8-bit grayscale versions used by our initial models to focus on colony morphology and texture. Using the gray-level transformation helps reduce sensitivity to background color and lighting artifacts that do not carry biologically meaningful information.

**Figure 6.**
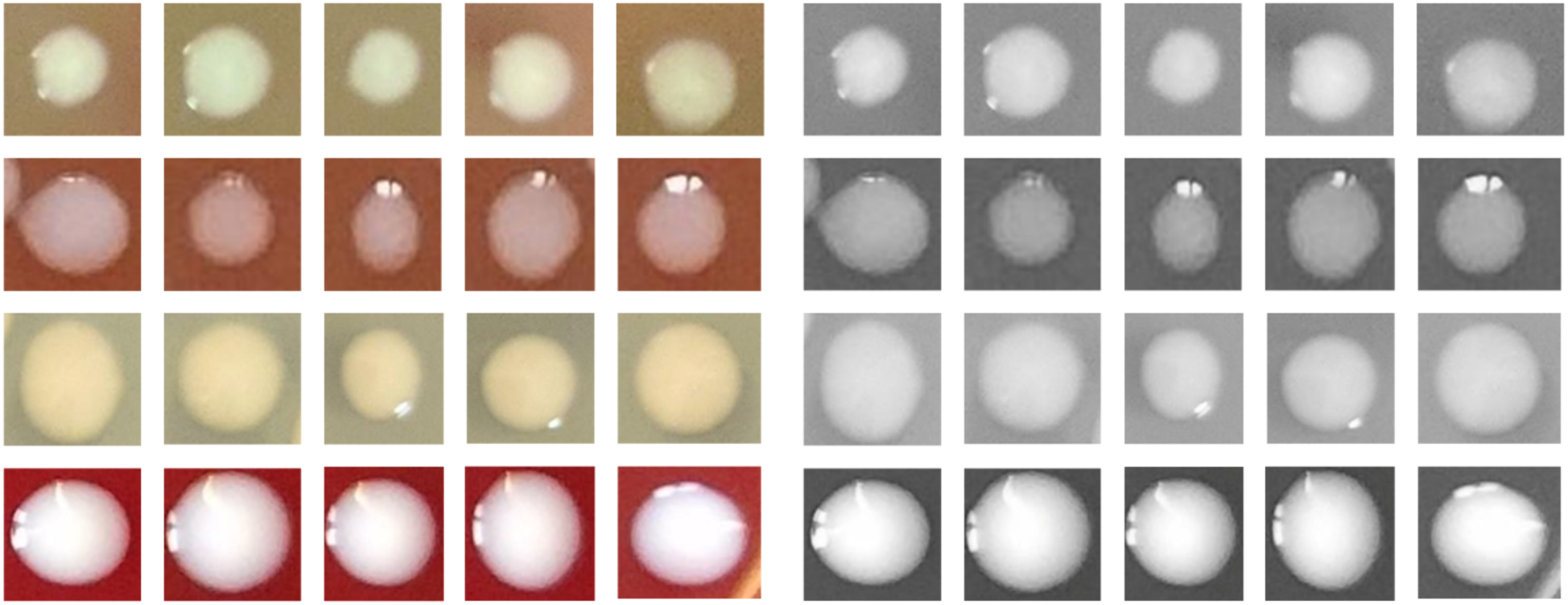
Grid of sample patches for the four public-dataset classes used in fine-tuning, arranged in 4 rows (one per class: sp14 *Listeria monocytogenes*, sp16 *Pasteurella multocida*, sp20 *Salmonella enterica*, sp22 *Staphylococcus hyicus*) and 5 columns (five representative colonies). Each cell shows the original color crop on the left and the corresponding 8-bit grayscale version on the right.

**Figure 7.**
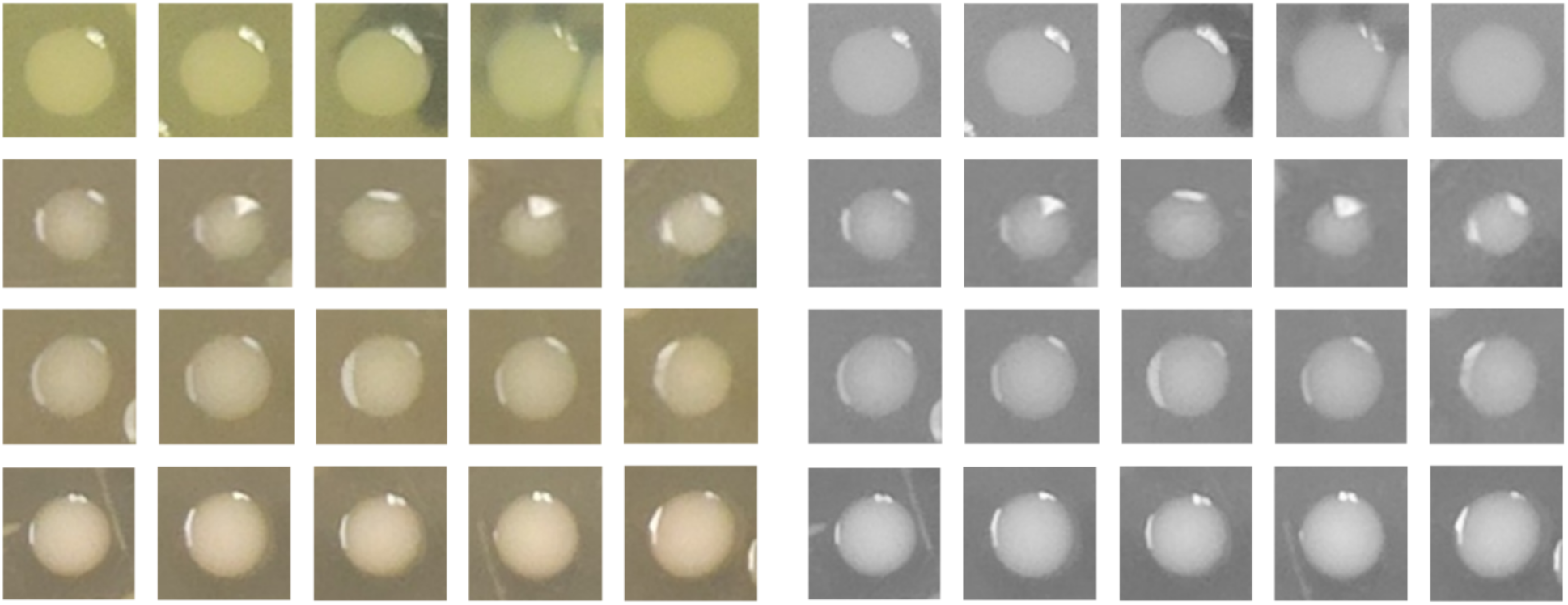
Representative detections for the four coliform species in the in-house (Target) dataset. From left to right: *Escherichia coli*, *Citrobacter freundii*, *Enterobacter aerogenes*, and *Klebsiella pneumoniae*. As in Figures 5 and 6, each pair shows the original color image (left) and the 8-bit grayscale version (right).

Given the small size and high density of many colonies, each full-plate image was subdivided into overlapping tiles (both 640 × 640 px and 1,280 × 1,280 px grids with 100 px and 200 px overlaps). This tiling strategy ensures that colonies near tile borders retain sufficient context for reliable detection. When using overlapping tiles, it’s common for the same colony to be detected multiple times near tile edges. A standard approach to address this redundancy is the use of Non-Maximum Suppression (NMS), a post-processing technique that filters overlapping predictions by retaining only the one with the highest confidence score based on the Intersection over Union (IoU) metric. Integrating NMS is especially useful in tiled object detection pipelines, as it can improve output clarity by consolidating redundant detections. While our current pipeline proceeds directly to classification using all predicted regions without applying NMS, this design choice does not substantially impact performance, as each detected patch is independently cropped, resized, and classified. Nonetheless, incorporating NMS in future iterations may help streamline detection outputs and reduce post-processing effort, particularly in densely populated regions or at tile boundaries.

We optimized tile size and overlap parameters via grid search, using the Adam optimizer with standard YOLOv8 hyperparameters. After 50 epochs, the model generalized well to unseen data, achieving an mAP@50 of 87.6 % on a held-out set of 105 public images. The resulting detector automatically produces tight bounding boxes around each colony, which are subsequently cropped, zero-padded to preserve aspect ratio, and resized to 100 × 100 px for the classification stage.

#### 2.2.3 Stage 2 - Colony Classification with CNN

To address the specific challenges of bacterial colony classification with a limited sample size, we designed a CNN architecture tailored to the morphological characteristics of microbial growth patterns. The model was pretrained on a domain-relevant public dataset of bacterial colonies, compensating for the lack of widely available pretrained models suited to this specific task. This pretraining improved feature generalization and accelerated convergence for both standard CNN architectures (such as ResNet commonly used for large-scale natural image classification [16]) and shallow custom CNN models. Our overall approach also allows finer control over learned representations, aligning more closely with the visual distinctions relevant to colony classification. Once colonies had been localized, each cropped region entered a two-phase transfer learning [17] workflow. In the first phase, we trained a base CNN model on a curated 10-class subset of the public dataset. This model was designed to learn generalized visual features, such as shape, edge orientation, and texture, across a diverse set of bacterial colonies. To avoid task leakage and promote feature transferability, we intentionally excluded classes closely related to those in our target dataset.

As shown in Figure 8 in detail, the base CNN architecture consists of three convolutional blocks. The first block applies a 2D convolution with 32 filters (3×3 kernel), followed by a 2×2 max pooling layer to reduce spatial dimensions. The second block uses 64 filters (3×3) and another 2×2 pooling layer to capture more complex patterns. The third block includes 128 filters (3×3) without additional pooling, enabling deeper feature extraction. These convolutional layers are followed by a global average pooling layer, a fully connected dense layer with 64 ReLU-activated units, and a softmax output layer with 10 units for classification during the pretraining phase.

**Figure 8.**
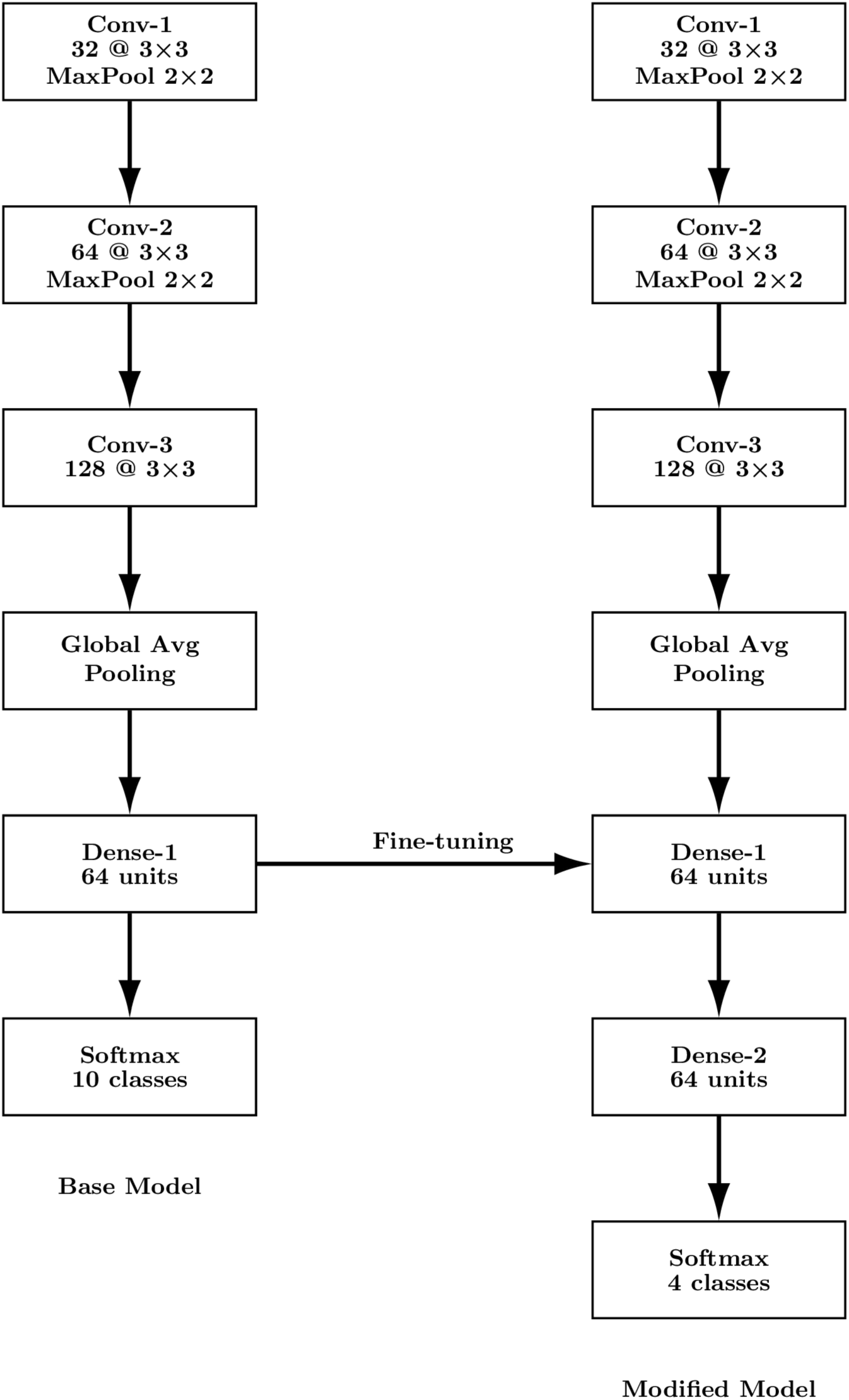
Transfer learning workflow for coliform colony classification. A base CNN is pretrained on a public bacterial colony dataset to learn generalizable morphological features. The learned convolutional layers are then reused and fine-tuned with a task-specific classification head using in-house coliform images, enabling robust performance with limited labeled data.

In the second phase, we transferred the learned weights from the base model to a new classification head tailored for our target task. Specifically, we removed the original softmax layer, appended a new dense layer (64 units, ReLU), and added a final softmax output layer with 4 units corresponding to the bacterial species in the in-house dataset. This fine-tuning was performed using colony patches extracted from the in-house high-resolution plate images.

The pretraining and fine-tuning workflow illustrated in Figure 8 on the lightweight custom CNN, is also extended to ResNet [16], a deeper pretrained architecture for color images. Specifically, we use ResNet-18 initialized with ImageNet-pretrained weights, which are first adapted using the intermediate 10-class bacterial classification task and subsequently fine-tuned on the target coliform dataset, following the same staged training procedure (this time using RGB images without converting them to gray).

#### 2.2.4. Training Procedure and Transfer Learning Performance Evaluation

In order to obtain the base CNN model shown in Figure 8, we “warmed up” the architecture by pretraining on ten taxonomically distinct classes drawn from the same 24-class public dataset (approximately 4,500 colony patches, 450 per class). We trained the model for 90 epochs using the Adam optimizer (learning rate = 1 × 10⁻⁴), with early stopping based on validation loss. This base-training phase yielded a mean validation accuracy of 77% on a 10-class colony classification task, evaluated on 500 test patches corresponding to those classes, which demonstrates that the network learned generalizable morphological features such as colony shape and texture (see Figures 4 and 8 for a schematic overview of the detection process and the usage of the base-task for the transfer learning process).

In the second phase, the pretrained CNN was fine-tuned separately on two four-class classification tasks: (i) For our in-house dataset comprising *Citrobacter freundii*, *Enterobacter aerogenes*, *Escherichia coli*, and *Klebsiella pneumoniae*, and (ii) a different four-class subset of the public dataset comprising *Listeria monocytogenes*, *Pasteurella multocida*, *Salmonella enterica*, *Staphylococcus hyicus*. As shown in Figure 8, the fine-tuning was performed by appending a fully connected Dense layer (Dense-2) on top of a frozen base CNN, without modifying the underlying architecture. We used an 80/20 train/test split and applied standard data augmentations (random horizontal/vertical flips and small-angle rotations) for the training phase. Inference time per plate-patch remained under five seconds on an AWS EC2 instance, demonstrating that the proposed transfer learning framework offers both high accuracy and practical efficiency for potential use in scalable, field-deployable water-quality monitoring systems.

### 2.3 Traditional Feature Extraction (Baseline Comparison)

To establish a performance baseline, we implemented three well-known, handcrafted feature descriptors including Histogram of Oriented Gradients (HOG) [18], Local Binary Patterns (LBP) [19], and Haralick texture features [20] and paired each with two classical classifiers (Support Vector Machine [21] and Random Forest [22]). By comparing these pipelines against our deep-learning approach, we quantify the gains afforded by end-to-end feature learning.

First, each 100×100-pixel colony patch was converted to an 8-bit grayscale image, matching the inputs used by our CNN. We then computed:

- **HOG:** Initially applied using default settings, 9 orientation bins, cell size of 8×8 pixels, and block size of 2×2 cells. We also performed hyperparameter tuning, which yielded optimal settings of: pixels_per_cell = (16, 16), cells_per_block = (2, 2), and orientations = 8.
- **LBP:** Used the default uniform Local Binary Pattern configuration with a radius of 1 pixel and 8 sampling points, producing 59-bin histograms. Varying the radius did not lead to improved performance.
- **Haralick:** Extracted 13 standard texture descriptors (e.g., contrast, correlation, energy, homogeneity) from gray-level co-occurrence matrices (GLCMs) computed at a distance of 1 pixel and angles of {0°, 45°, 90°, 135°}. Hyperparameter tuning did not yield performance gains.

All feature vectors were z-score standardized before classification. For each descriptor set, we trained:

- **Support Vector Machine (SVM):** with RBF kernel, cost parameter C and kernel width γ optimized via grid search over {10⁻², 10⁻¹, 1, 10} with five-fold cross-validation.
- **Random Forest (RF):** For all handcrafted feature sets, classification was performed using a Random Forest with 100 trees. We tuned the maximum depth over the set {None, 10, 20}, and at each split, √p features were randomly selected (where p is the number of input features).

## 3. Results

### 3.1 Colony Detection Accuracy with YOLOv8

We evaluated our YOLOv8-based colony detector on 105 held-out images drawn from the public dataset. The model achieved a precision of 0.906 and a recall of 0.827, corresponding to an mAP@50 of 0.877. When measured across the full IoU spectrum (mAP@50-95), the detector yielded 0.451, indicating robust localization under varying overlap thresholds. Compared to our initial ImageJ workflow that identified 185 colonies across the four in-house culture plates, the YOLO detector located 248 colonies on the same plates, demonstrating its ability to discover subtler instances that manual annotation missed. As before, we used an 80/20 train/test split, rounding the test set down to 49 examples rather than up to 50, in order to avoid overly round accuracy values and better reflect reporting granularity.

### 3.2 Improving Classification Accuracy with Transfer Learning

We evaluated classification performance under several regimes, starting with a manual workflow based on ImageJ. Using ImageJ-cropped “tight” bounding boxes from our four-class in-house dataset, a CNN trained from scratch achieved 73% accuracy. When trained instead on slightly larger, loosely cropped patches around each colony, accuracy improved to 83%, suggesting that overly tight crops may omit visual context important for robust classification.

Next, we incorporated transfer learning by pretraining the CNN on ten public colony classes and fine-tuning it on the in-house dataset, which further boosted accuracy to 94% (±3.6% over ten runs). However, this result may appear deceptively strong, as it was based on a manually curated subset; ImageJ failed to detect as many colonies as YOLOv8, missing a number of subtler or overlapping instances. Thus, the effective classification task involved fewer and more distinct examples. To address this limitation and improve scalability, we replaced the manual annotation step with automated colony detection using YOLOv8. This YOLO+CNN pipeline processes raw plate images end-to-end—without the need for ImageJ and enables consistent classification of all visible colonies, including small and densely clustered ones.

To assess the end-to-end pipeline (YOLO + CNN) on the in-house plates, we first used automatic cropping via our trained YOLO (Figure 4) and classification by the transfer-learned CNN (Figure 8). This proposed pipeline yielded an average accuracy of ≈ 86%, with the best-performing model achieving ≈ 92% on held-out in-house patches.

Finally, to assess generality (applicability of our method to other small datasets), we applied our pipeline to an independent four-class subset of the original 24-class public dataset. This subset comprised two classes from our initial ten (with different, randomly selected plate images) plus two novel classes drawn from the remaining 14, for a total of 17 agar-plate images. YOLO detected 1,791 colonies across those plates, and the CNN achieved 92% classification accuracy on the corresponding test split. Together, these results underscore the usefulness of transfer learning in accelerating convergence and substantially improving accuracy across both in-house and larger public-dataset tasks.

Tables 1 and 2 present a direct comparison of our traditional ML baselines and the proposed CNN pipeline, using a uniform header in each: Model, Mean Accuracy (%), Std Dev (%), Min Accuracy (%), and Max Accuracy (%), which are computed from 10 independent runs (using different train-test splits). Table 1 reports these classification statistics on the Target Dataset (in-house four coliform classes: *Citrobacter freundii, Enterobacter aerogenes, Escherichia coli, Klebsiella pneumoniae*). Table 2 shows analogous experiments on the 4 classes from the big public dataset: *Listeria monocytogenes, Pasteurella multocida, Salmonella enterica, Staphylococcus hyicus*). In both tables, entries labeled “Optimized” denote feature-classifier pipelines whose hyperparameters were exhaustively tuned with scikit-learn’s GridSearchCV (cross-validating over SVM kernels and regularization strengths, Random Forest tree counts, etc.). All other rows correspond to default-parameter settings. Systematic tuning via GridSearchCV consistently produced higher test accuracies and tighter standard deviations of those accuracies across all the runs, which demonstrate improved validation performance and generalization over the out-of-the-box configurations.

**Table 1.** Experimental results on the target dataset (in-house dataset with 4-classes [13]), used for fine-tuning. The dataset includes four bacterial classes: *Citrobacter freundii, Enterobacter aerogenes, Escherichia coli, and Klebsiella pneumoniae*.

| Model | Mean Accuracy (%) | Std Dev (%) | Min Accuracy (%) | Max Accuracy (%) |
| --- | --- | --- | --- | --- |
| HOG + SVM | 76.73 | 5.78 | 69.38 | 89.79 |
| HOG + RF | 73.26 | 6.48 | 63.26 | 87.75 |
| HOG + SVM (optimized) | 83.87 | 3.81 | 77.55 | 91.83 |
| HOG + RF (optimized) | 80.20 | 3.65 | 75.51 | 87.75 |
| LBP + SVM | 60.61 | 5.40 | 53.06 | 71.42 |
| LBP + RF | 66.53 | 5.71 | 57.14 | 75.51 |
| Haralick + SVM | 65.31 | 4.47 | 59.18 | 75.51 |
| Haralick + RF | 79.59 | 5.77 | 69.38 | 87.75 |
| Proposed Method | 85.91 | 4.12 | 79.59 | 91.83 |

**Table 2.**
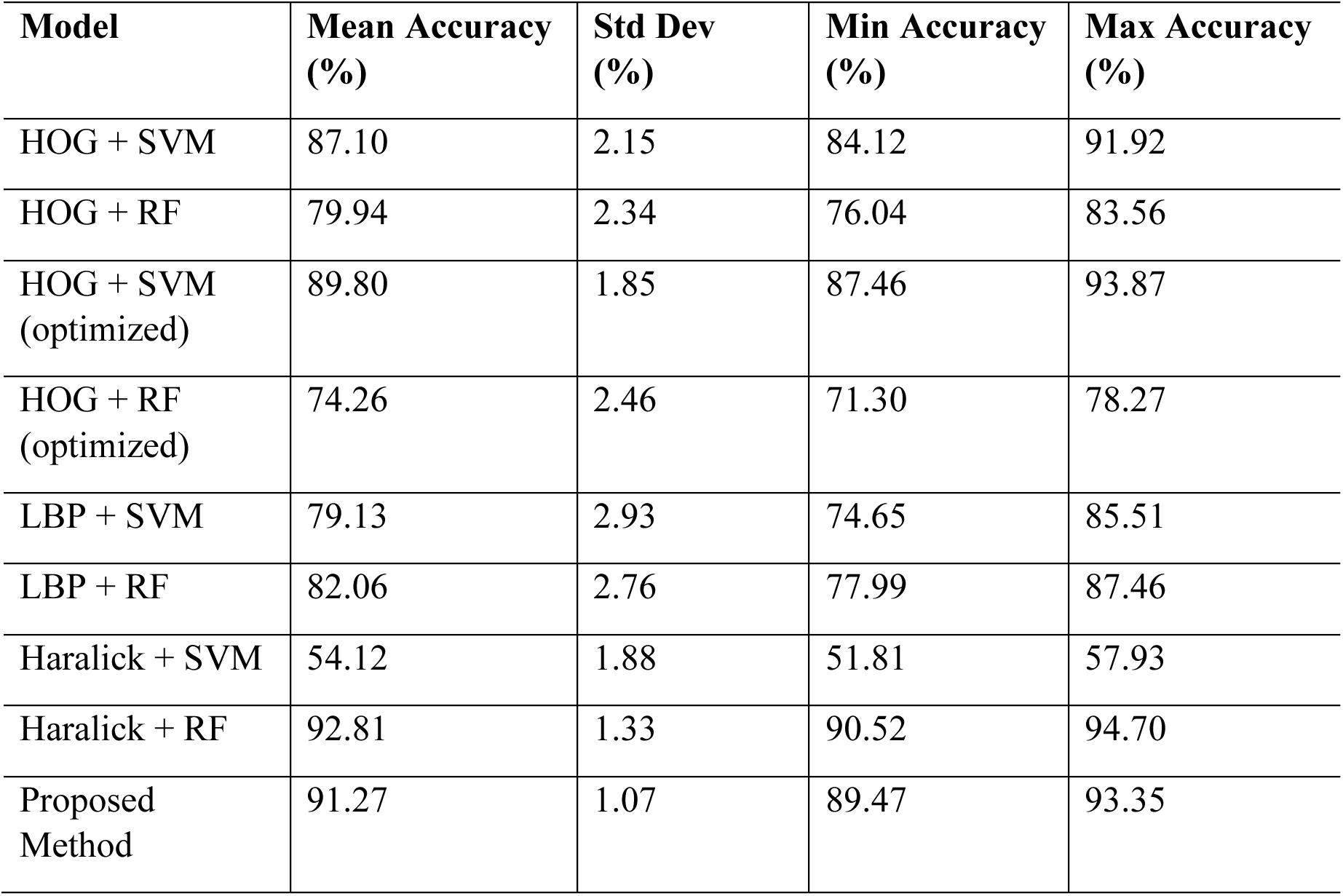
Classification results on a four-class subset of the public bacterial colony dataset [12], which also served as the pretraining base for transfer learning. The four bacterial classes are: *Listeria monocytogenes, Pasteurella multocida, Salmonella enterica, and Staphylococcus hyicus*.

| Model | Mean Accuracy (%) | Std Dev (%) | Min Accuracy (%) | Max Accuracy (%) |
| --- | --- | --- | --- | --- |
| HOG + SVM | 87.10 | 2.15 | 84.12 | 91.92 |
| HOG + RF | 79.94 | 2.34 | 76.04 | 83.56 |
| HOG + SVM (optimized) | 89.80 | 1.85 | 87.46 | 93.87 |
| HOG + RF (optimized) | 74.26 | 2.46 | 71.30 | 78.27 |
| LBP + SVM | 79.13 | 2.93 | 74.65 | 85.51 |
| LBP + RF | 82.06 | 2.76 | 77.99 | 87.46 |
| Haralick + SVM | 54.12 | 1.88 | 51.81 | 57.93 |
| Haralick + RF | 92.81 | 1.33 | 90.52 | 94.70 |
| Proposed Method | 91.27 | 1.07 | 89.47 | 93.35 |

As shown in Table 1, on our 4-class in-house data test split, the best baseline was HOG + SVM (≈ 84% accuracy), followed by HOG + RF (≈ 80%) and Haralick + RF (≈ 80%), all of which fell short of the ≈ 86% achieved by our CNN fine-tuned with transfer learning. These results underscore the superiority of learned deep features over handcrafted descriptors for robust, high-throughput bacterial-colony classification. Similarly, as shown in Table 2, the proposed method remained among the top-performing approaches, with only a slight edge in accuracy observed for Haralick + RF, a non-transfer-learning baseline. The comparatively lower performance of Haralick + RF on the in-house dataset (Table 1) underscores the consistency and generalizability of our transfer learning approach.

### 3.3 Evaluation of Classification Performance on Colored Colony Images

To assess the sensitivity of the proposed pipeline to color information and architectural choices, we conducted additional classification experiments using RGB colony images and a ResNet-18 backbone in place of the earlier shallow CNN. These experiments are not intended to replace the grayscale analysis, but rather to evaluate how incorporating color cues and a deeper, widely used architecture affects performance and generalization.

For this analysis, the CNN was initialized via transfer learning using a slightly different set of ten bacterial species from the public dataset (sp02, sp05, sp06, sp07, sp10, sp14, sp16, sp19, sp21, sp23), selected to span a range of colony morphologies while ensuring sufficient sample availability. A total of 1,000 colored colony patches were used per class, with 800 samples for training and 200 for testing the base CNN. Pretraining was performed on these patches, after which the model was fine-tuned for downstream 4-class classification tasks.

For the pretraining phase, the classifier achieved strong and balanced performance, with precision, recall, and F1-score all exceeding 0.92 and an overall accuracy of approximately 92.5% (Table 3 and Figure 9). To further assess feature transferability, the pretrained base network was evaluated on an independent four-class subset of the public dataset (sp05, sp10, sp22, sp13), where it achieved an accuracy of 96.4% (Table 4 and Figure 10). When transferred and fine-tuned on the in-house dataset, the model achieved the same level of high accuracy (Table 5 and Figure 10). To further assess robustness under limited data, k-fold cross-validation (k = 10) was performed on the in-house dataset. The model achieved a mean accuracy of 98.0% with a standard deviation of 2.8%. Macro-averaged precision, recall, and F1-score were 98.4% ± 2.3%, 97.8% ± 3.4%, and 97.8% ± 3.2%, respectively. These results indicate stable performance across folds and suggest limited sensitivity to the specific train/test partition within a given experimental setting.

**Figure 9.**
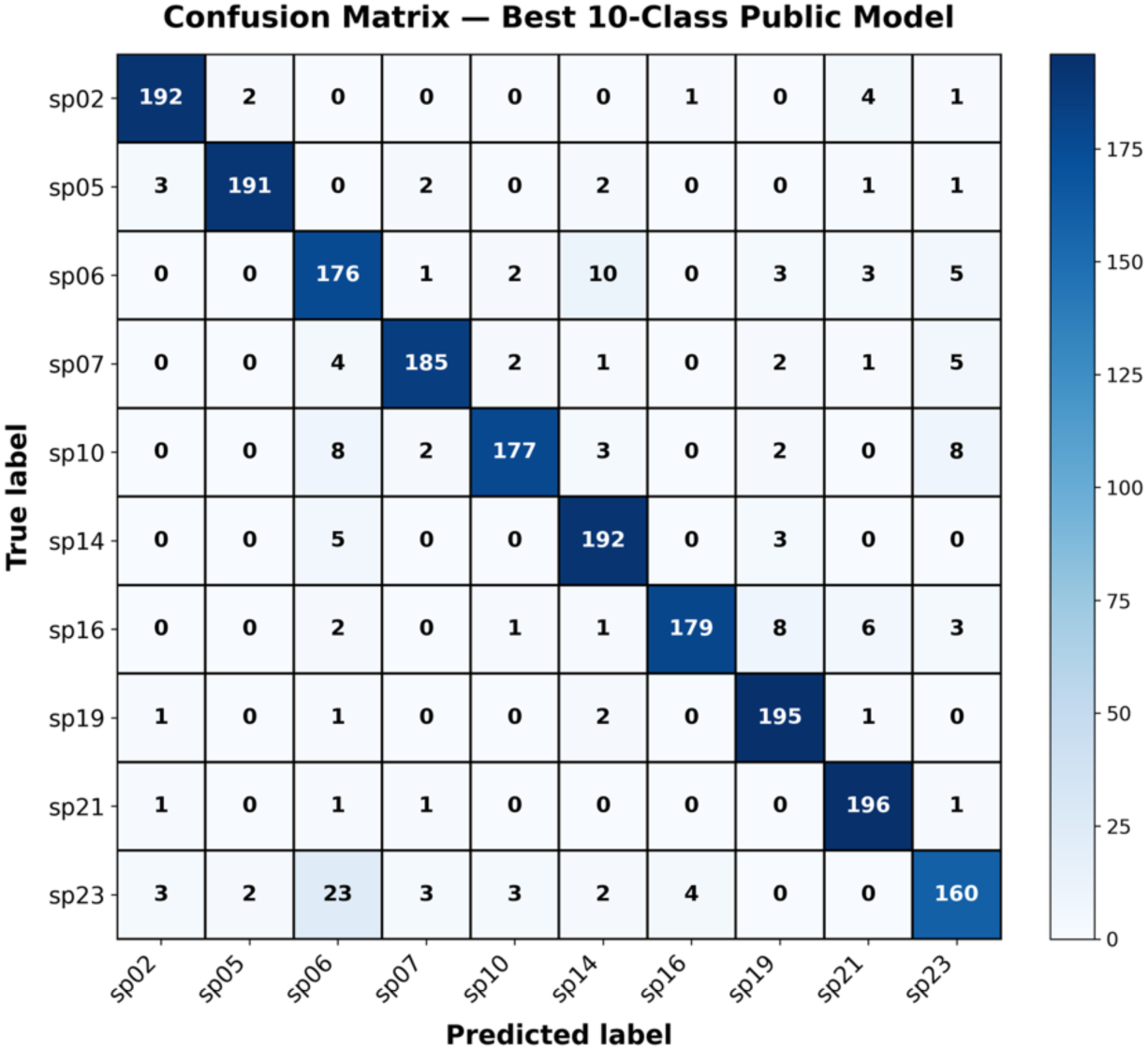
Confusion matrix of a sample run for the public 10-class dataset. Darker blue cells indicate more correct predictions, while lighter cells reflect fewer or misclassified instances.

**Figure 10.**
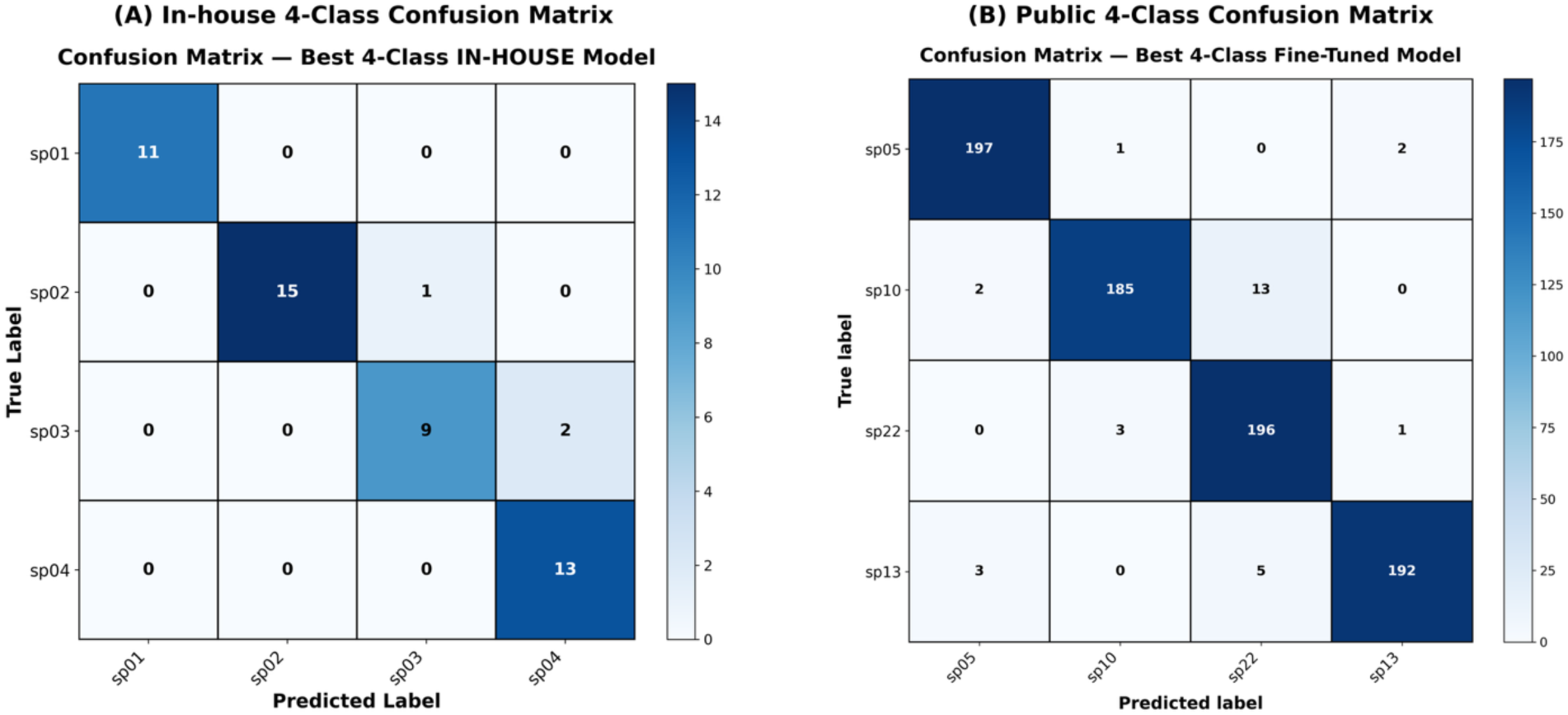
Subplot A shows the confusion matrix of a sample run for the in-house 4-class classification task. Subplot B shows the confusion matrix for the public 4-class dataset.

**Table 3.** Per-class precision, recall, F1-score, and support for the 10-class public bacterial colony dataset using the ResNet-18–based classifier with RGB inputs.

| Class | Precision | Recall | F1-score | Support |
| --- | --- | --- | --- | --- |
| sp02 | 0.96 | 0.96 | 0.96 | 200 |
| sp05 | 0.979 | 0.955 | 0.967 | 200 |
| sp06 | 0.8 | 0.88 | 0.838 | 200 |
| sp07 | 0.954 | 0.925 | 0.939 | 200 |
| sp10 | 0.957 | 0.885 | 0.919 | 200 |
| sp14 | 0.901 | 0.96 | 0.93 | 200 |
| sp16 | 0.973 | 0.895 | 0.932 | 200 |
| sp19 | 0.915 | 0.975 | 0.944 | 200 |
| sp21 | 0.925 | 0.98 | 0.951 | 200 |
| sp23 | 0.87 | 0.8 | 0.833 | 200 |
| Accuracy |  |  | 0.921 | 2000 |
| Macro Avg | 0.923 | 0.922 | 0.921 | 2000 |
| Weighted Avg | 0.923 | 0.921 | 0.921 | 2000 |

**Table 4.** Per-class classification performance for the four-class public bacterial colony dataset using a ResNet-18 classifier pretrained via transfer learning on RGB colony images.

| **Class** | **Precision** | **Recall** | **F1-score** | **Support** |
| --- | --- | --- | --- | --- |
| sp05 | 0.975 | 0.985 | 0.98 | 200 |
| sp10 | 0.979 | 0.925 | 0.951 | 200 |
| sp22 | 0.916 | 0.98 | 0.947 | 200 |
| sp13 | 0.985 | 0.96 | 0.972 | 200 |
| **Accuracy** |  |  | 0.963 | 800 |
| **Macro Avg** | 0.964 | 0.963 | 0.963 | 800 |
| **Weighted Avg** | 0.964 | 0.963 | 0.963 | 800 |

**Table 5.** Per-class classification performance for the four-class in-house bacterial colony dataset using a ResNet-18 classifier pretrained via transfer learning on RGB images from the public dataset.

| **Class** | **Precision** | **Recall** | **F1-score** | **Support** |
| --- | --- | --- | --- | --- |
| sp05 | 1 | 1 | 1 | 11 |
| sp10 | 1 | 0.938 | 0.968 | 16 |
| sp22 | 0.917 | 1 | 0.957 | 11 |
| sp13 | 1 | 1 | 1 | 13 |
| **Accuracy** |  |  | 0.98 | 51 |
| **Macro Avg** | 0.979 | 0.984 | 0.981 | 51 |
| **Weighted Avg** | 0.982 | 0.98 | 0.981 | 51 |

## 4. Discussion

Our AI-powered pipeline achieved dramatic reductions in analysis time, shortening colony detection and classification from hours of manual work to under five seconds per plate. Transfer learning was especially impactful: by “warming up” the CNN on a diverse, 10-class public dataset, we minimized the amount of in-house -specific data required to reach high accuracy. With the transfer learning approach, the proposed model accelerated model convergence and also mitigated overfitting on our limited in-house dataset.

Comparing our YOLOv8-based detection to the original ImageJ workflow underscores the strength of deep learning for localization: YOLO detected 248 colonies on the four in-house culture plates, which is 35% more than the 185 colonies annotated using the manual ImageJ process. This automation helps deliver end-to-end and real-time inference. Additionally, these gains reflect both YOLO’s capacity to spot subtle, low-contrast colonies and its robustness to plate artifacts. In the classification stage, deep features learned by our transfer-learned CNN consistently ranked among the top-performing methods. As an end-to-end neural network, it offers greater flexibility and generalizability that enables the integration of additional features such as color and contextual metadata. These advantages allow the CNN-based approach to outperform traditional pipelines built on handcrafted descriptors such as HOG, LBP, and Haralick.

In addition to advancing technical capabilities for microbial monitoring, this study presents a substantially reproducible, low-cost pipeline that combines wet-lab culturing, smartphone imaging, and deep learning for colony detection and classification. The data-collection and annotation workflows were intentionally designed for accessibility in educational and interdisciplinary research environments, enabling students and non-CS collaborators to contribute meaningfully without specialized equipment.

Taken together, the experimental results support the stability of the proposed two-stage framework across different input modalities and model configurations. While color information can provide additional discriminative cues for certain species, overall performance trends remain consistent with those observed under grayscale preprocessing. This framework therefore supports both scalable field deployment and straightforward adoption in resource-limited settings. However, we observe that cross-plate generalization without prior adaptation remains challenging, underscoring the importance of limited task-specific fine-tuning (warm-up) in practical deployments: Figure 11 illustrates the 80/20 train/test split on plate-level data for the in-house dataset and the limited nature of the spatial separation of training and testing samples on each plate. Training patches are indicated by blue bounding boxes, while test-set predictions are shown in green for true positives and yellow for false negatives.

**Figure 11.**
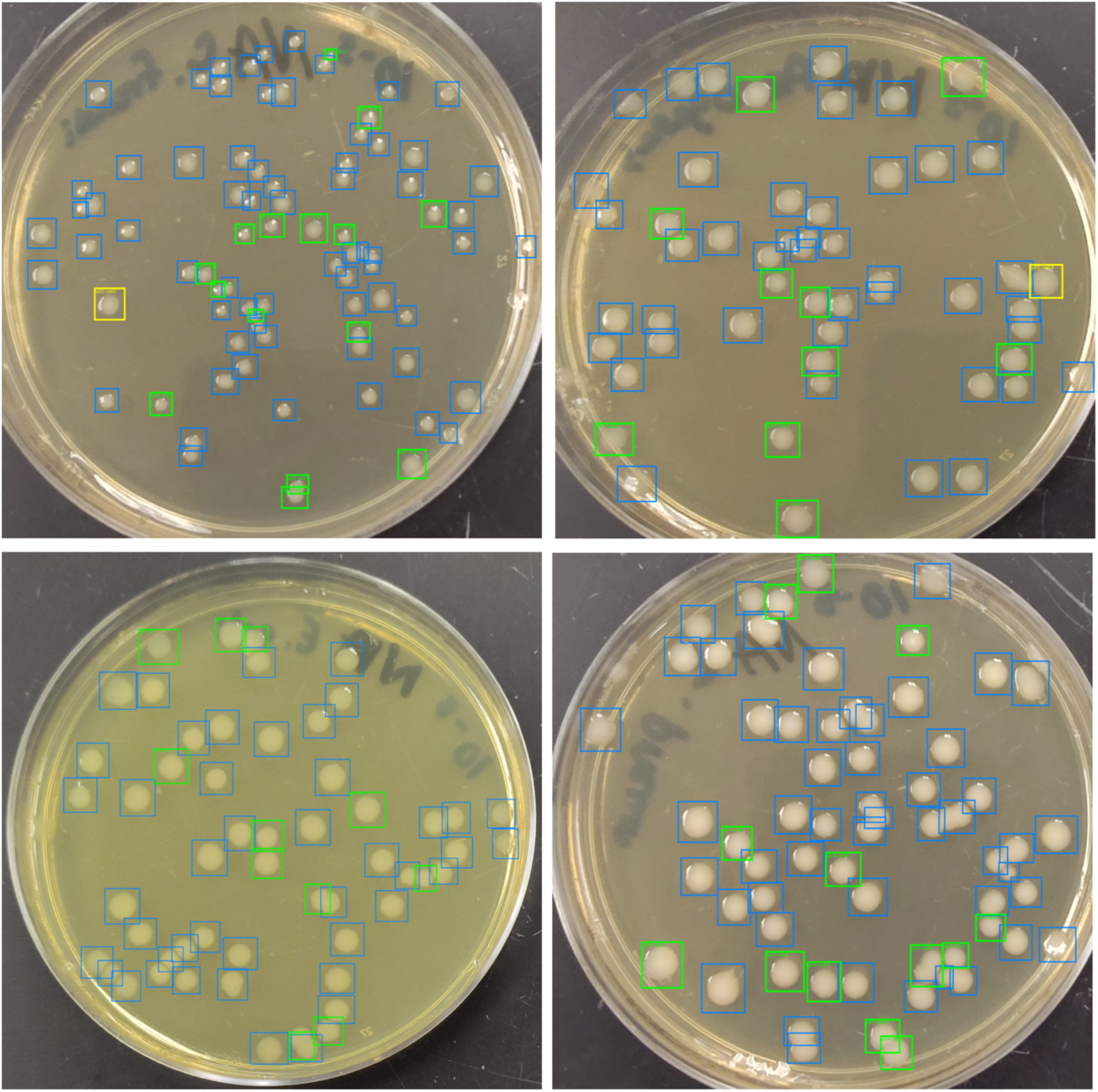
Visualization of plate-level training and testing splits for the in-house four-class dataset under an 80-20 protocol. Training patches are shown with blue bounding boxes. During testing, correctly classified colonies are shown in green, while missed detections (false negatives) are shown in yellow.

## 5. Conclusions

Our pipeline demonstrated that combining YOLOv8 for object detection with a transfer-learned CNN can deliver accurate and rapid bacterial colony identification and classification, with potential application to water quality monitoring. By reducing processing time to under five seconds per plate and achieving over 90% accuracy, the system is well suited for scalable and field-deployable applications. However, further fine tuning of the pipeline is needed to improve detection accuracy. To extend both the scientific reach and practical utility of our pipeline, we have identified the following complementary directions. Future work will focus on improving robustness and generalization by incorporating color imagery and relevant environmental metadata, as well as exploring self-supervised learning to reduce labeling burden in settings where mixed-culture plates are abundant. In parallel, we aim to translate the system into accessible web-based interfaces and integrate it into broader AI-driven educational and decision-support frameworks.

## CRediT authorship contribution statement

**Shivaji Mallela**: Writing - review & editing, Writing -- original draft, Software, Visualization, Validation; **Abria Gates**: Writing - review & editing, Writing -- original draft, Software; **Sandeep Medepalli**: Software; Data curation; Validation; Writing - review; **Benedict Okeke**: Writing - review & editing, Writing -- original draft, Visualization, Validation, Funding acquisition, Data curation; **Olcay Kursun**: Writing - review & editing, Writing -- original draft, Visualization, Validation, Supervision, Software, Resources, Project administration, Methodology, Investigation, Funding acquisition, Formal analysis, Data curation, Conceptualization.

## Acknowledgements

This work was supported by the NSF grant 2435093 (ExpandAI) and the NSF LSAMP Award # 1712692 - The Greater Alabama Black Belt Region (GABBR) LSAMP. The authors acknowledge the early involvement of Dr. Sutanu Bhattacharya in the broader project context supported by the NSF ExpandAI initiative.

## Data Availability

The public dataset used in this study is available at: https://doi.org/10.1038/s41597-023-02404-8. The complete in-house dataset and annotation guidelines are available on GitHub [13] and also from the authors upon request.

